# Genotype dependence of virulence-associated genes in ESBL-producing *Escherichia coli* isolated from the blood of patients with urinary tract infections

**DOI:** 10.1101/2024.08.21.608955

**Authors:** Mayuko Tanaka, Tomoko Hanawa, Tomoya Suda, Yasunori Tanji, Le Nhat Minh, Kohei Kondo, Aa Haeruman Azam, Kotaro Kiga, Shota Yonetani, Ryu Yashiro, Takuya Omori, Takeaki Matsuda

**Affiliations:** Department of General Medicine, Kyorin University School of Medicine, Tokyo, 181-8611, Japan; Department of Traumatology and Critical Care Medicine, Kyorin University School of Medicine, Tokyo, 181-8611, Japan; School of Life Science and Technology, Tokyo Institute of Technology, Nagatsutacho, 226- 8501, Yokohama, Japan; Antimicrobial Resistance Research Center, National Institute of Infectious Disease, Tokyo, 189-0002, Japan; Research Center for Drug and Vaccine Development, National Institute of Infectious Diseases, Tokyo 162-8640, Japan; Department of Medical Technology, Faculty of Health Sciences, Kyorin University, Tokyo, 181-8612, Japan; Center for Data Science Education & Research, Kyorin University, Tokyo, 181-8612, Japan

## Abstract

Approximately 80% of urinary tract infections (UTIs) are caused by uropathogenic *Escherichia coli* (UPEC). The prevalence of extended-spectrum β-lactamase (ESBL)- producing *E. coli* in UPEC isolates belonging to specific multilocus sequence-type clones is a global concern. Relations between the pathogenicity of UPEC virulence factors and genotype have been discussed. However, the specific virulence factors and genotypes associated with a higher likelihood of causing UTIs remain unclear. This study analyzed the genotypes and virulence-associated genes of 46 ESBL-producing strains isolated from the blood of patients with UTIs who visited the Department of Emergency Medicine from 2017 to 2022 to elucidate the characteristics of UPEC strains associated with severe infections. Most phylogroups of clinical isolates were B2, except for D, found in three strains. The dominant multilocus sequence typing (MLST) was ST131, followed by ST73, ST95, and ST38, frequently observed in UPEC strains. ST131 strains were more resistant to levofloxacin (87.5%) than non-ST131 strains (50.0%) and caused fewer sepsis cases than non-ST131 strains. Based on *in silico* analysis, of 23 clinical isolates, the genes detected in all clinical strains could have important role in invasive UTIs. Clustering analysis highlighted the genotype MLST dependence of UPEC-specific virulence-associated genes. The absence of several UPEC-specific gene loci or the presence of ST38-specific indicated atypical repertories of virulence-associated genes in ST38 strains. Genes encoding secretion systems, which were found in enteropathogenic *E. coli*, were less detected in ST131 strains. These results suggested that the correlation of MLST and repertories should be considered to understand UPEC virulence.

**IMPORTANCE STATEMENT:** Several virulence genes, their diverse repertories depending on the strains, and various pathological indications complicate the pathogenicity of uropathogenic *Escherichia coli* (UPEC). The genes detected in all clinical isolates in this study would play a crucial role in UPEC pathogenicity. In contrast, depending on the genotype, the distribution of UPEC- specific genes and other pathogenic *E. coli* virulence genes in some genotype strains varied. Secretion system genes related to the pathogenicity of enteropathogenic *E. coli* (EPEC) were less in the strains of the most prevalent genotype, ST131; this was relevant to the number of sepsis cases. These results clearly indicated genotype dependencies on virulence-associated gene repertories and ambiguity in UPEC and EPEC. This study proposed that analyzing genotype-specific virulence factors and sharing virulence genes with EPEC will bring new insights into UPEC pathogenicity.

## INTRODUCTION

Pathogenic *Escherichia coli* is classified as enteropathogenic *E. coli* (EPEC), which causes diarrhea, and extraintestinal pathogenic *E. coli* (ExPEC), which causes urinary tract infections (UTIs), bacteremia, or sepsis (1). Extended-spectrum β-lactamase (ESBL)- producing *E. coli*, one of the critical multidrug-resistant bacteria, is disseminated in the community and healthcare facilities and the environment, representing public health concerns (2 ,3). Most ExPEC belong to phylogroup B2, followed by D, among the seven classified phylogroups (A, B1, B2, and D–G) (4). Contrary to the varied multilocus sequence typing (MLST) of ESBL-producing *E. coli* detected, most strains isolated from human bloodstream infections or UTIs belong to ST131 of phylogroup B2 and other limited sequence types worldwide (5, 6, 7, 8).

UTIs are detected in all age groups and are prevalent in women, who will experience UTIs at least once in their lives (9). Uropathogenic *E. coli* (UPEC) involves 80% of UTIs (10). UPEC indicates diverse pathology from noninvasive UTIs, such as asymptomatic bacteriuria cystitis, to invasive UTIs, such as pyelonephritis, possibly presenting bacteremia and sepsis. Bacterial virulence strongly affects the clinical course of UTI, besides host immunity. Therefore, UTIs can develop into invasive infections, life-threatening diseases, pyelonephritis, and sepsis (urosepsis), whereas UPEC represents common diseases. These complications make the features of strains unclear.

Virulence-related genes that characterize UPEC and have UPEC-specific virulence genes, such as colonization factors, iron acquisition systems, and toxins, are frequently contained in pathogenicity islands found in UPEC (10, 11, 12). Although virulence-associated genes have been studied using many UPEC strains, mainly by polymerase chain reaction (PCR), the repertoire of virulence genes possessed by *E. coli* is diverse, and their roles in the pathogenesis remain unclear (13).

*E. coli* is the leading pathogen for global deaths (counts) attributable to and associated with bacterial antimicrobial resistance in 2019, and these diseases were due to extraintestinal infections (14). It is helpful to understand ESBL-producing UPEC causing invasive infections ought to predict the clinical pathway depending on the strains. However, bacterial characteristics that define severe invasive infections have not yet been clarified.

In this study, the genotype and virulence-associated genes of clinical isolates from blood with UTIs caused by ESBL-producing *E. coli* were characterized by PCR-based analysis of 46 strains and whole *in silico* analysis of genome sequencing using 23 isolates of non-ST131 and ST131 strains. Results revealed a dependence on MLST among UPEC-specific genes or gene locus on possessing virulence-associated genes. At the same time, some adhesins, iron acquisition systems, and serum resistance factors were present in all strains of clinical isolates. Based on the number of sepsis cases and virulence-associated genes in the genomes, the ST131 clone has less pathogenicity than the non-ST131 strains among UPEC causing bloodstream infections.

## MATERIALS AND METHODS

### Cases of ESBL-producing *E. coli* infections and clinical isolates

Sixty ESBL-producing *E. coli* strains isolated from blood cultures in routine clinical practice at the Departments of Emergency Medicine and Emergency General Medicine, Kyorin University Hospital, from 2017 to 2022, were included. The same strain from the same patient was considered the first isolate, and duplicates were deleted. The primary source of infection and genotype of bacterial strains are shown in Table 1.

**Table 1.** Primary source of infection and genotype of bacterial strains.

| **Primary source of infection** | **Total**<br>**(n=60) (%)** | **B2**<br>**(n=53)** | **D**<br>**(n=6)** | **F**<br>**(n=1)** |
| --- | --- | --- | --- | --- |
| Urinary tract infection | 46 (76.7) | 43 | 3 | 0 |
| Biliary tract infection | 9 (15.0) | 7 | 1 | 1 |
| Others | 5 (8.3) | 3 | 2 | 0 |

In cases where bacteria detected in urine, bile, or ascites cultures matched those identified in blood cultures, the respective organ was identified as the source of infection. In cases where only blood cultures were performed, the organ with apparent infection was considered the primary site. Cases in which the infected organ could not be identified were classified as unexplained were included in others together with gastrointestinal perforation. Data are expressed as number or percent (%) of the strains.

BD Phoenix (Becton Dickinson, Japan) was used for identification and drug susceptibility testing. Drug susceptibility testing was performed using the Clinical and Laboratory Standards Institute (CLSI)-compliant microliquid dilution method to determine minimum growth inhibitory concentrations for 20 antibiotics, and strains meeting the CLSI screening criteria for ESBL production were selected for confirmation as defined by the CLSI. ESBL production was determined using the double-disk synergy test, a confirmation test established by the CLSI.

Bacteremia was defined as the detection of ESBL-producing *E. coli* in blood culture. Sepsis was determined according to clinical criteria: suspected or proven focus of infection and acute elevation of Sequential Organ Failure Assessment score ≥2 points (a proxy for organ dysfunction) according to the Japanese Clinical Practice Guidelines for Management of Sepsis and Septic Shock 2016 and 2020.

### Genotyping by PCR and analysis of virulence-associated gene retention PCR analysis

DNA was extracted according to the Wizard® Genomic DNA Purification Kit (Promega) instructions and used for PCR. Phylogroup was determined according to Clermont’s scheme (15, 16). The representative virulence-associated genes (*papAH*, *papGII*, *papGIII*, *cnf1*, *hlyA*, *kpsMTII*, *fyuA*, *iutA*, *usp*, *malX*, *traT*, *ompT*, *fimH*, and *csgA*) associated with UPEC were detected by PCR. PCR was performed using the primers shown in Table 2. ST131 genotype was determined using the CicaGeneus® *E. coli* POT kit (Kanto Chemical Co., Ltd.) according to the instructions, and negative results were considered non-ST131.

**Table 2.**
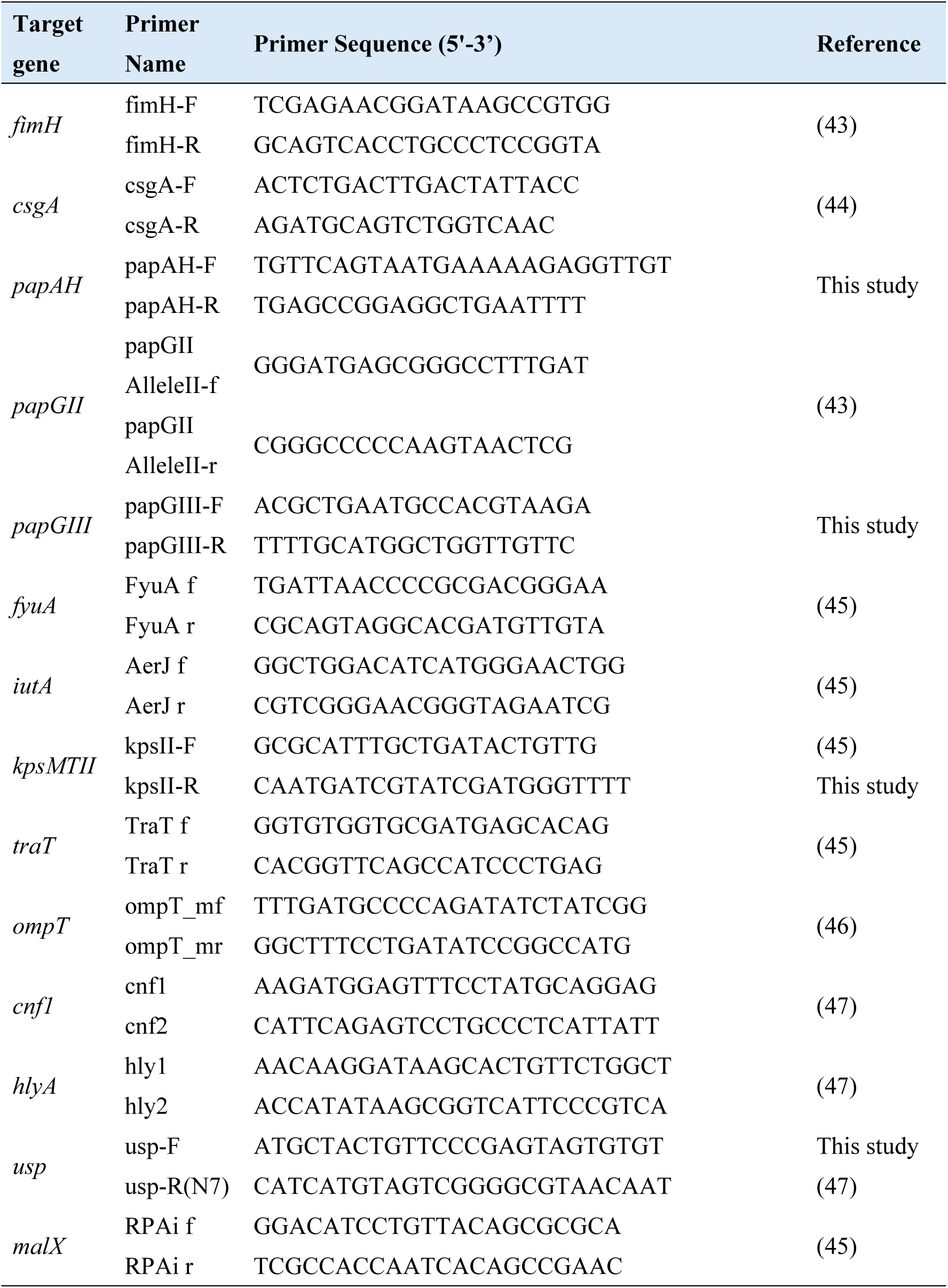
PCR primers used in this study.

### Whole-genome sequencing and genotyping *de novo* assembly and annotation

Clinical isolates were cultured in LB broth at 37°C for 15 h, and entire genomes were extracted using the Wizard® HMW DNA Extraction Kit (Promega). Library preparation was performed using the QIAseq FX DNA Library Kit (Qiagen, Hilden, Germany). Paired-end sequencing was performed on the DNBSEQ platform. Sequence reads were assembled *de novo* to contigs using Shovill version 1.1.0 (https://github.com/tseemann/shovill). Annotation was performed using Prokka version 1.14.6 (https://github.com/tseemann/prokka). Genomic information for the 23 strains was deposited in GenBank (BioProject ID PRJDB18240).

### In silico analysis

Phylogroup, MLST, serotype, and fimH and FumC type were analyzed by ClermonTyping (http://clermontyping.iame-research.center), mlst version 2.23.0 (https://github.com/tseemann/mlst), SerotypeFinder 2.0, and CHTyper 1.0 (Center for Genomic Epidemiology; https://cge.food.dtu.dk/services/SerotypeFinder/, https://cge.food.dtu.dk/services/CHTyper/), respectively.

ABRicate version 1.0.1 (https://github.com/tseemann/abricate) was used to detect virulence-associated genes, antibiotic resistance genes, and plasmids using the default parameters. *E. coli*_VF (https://github.com/phac-nml/Ecoli_vf), Resfinder (17, 18), and National Center for Biotechnology Information (NCBI) AMRFinderPlus (18) were used as databases. Basic Local Alignment Search Tool (BLAST) search of proteins or DNA sequences was performed at the NCBI (https://blast.ncbi.nlm.nih.gov/Blast.cgi?PROGRAM=blastp&PAGE_TYPE=BlastSearch&BLAST_SPEC=&LINK_LOC=blasttab&LAST_PAGE=blastn) to predict the gene functions.

### Clustering analysis

Clustering analysis heatmap and dendrogram generation were performed using Seaborn version 0.12.02 (statistical data visualization; https://seaborn.pydata.org). Reference strains for ESBL-producing ST131 UPEC were EC598 (GenBank accession no. HG941718), CFT073 (GenBank accession no. AE014075.1), UTI89 (GenBank accession no. CP000243), and 536 (GenBank accession no. CP000247) genome sequences were used for analysis. Other genomes with UPEC strains of phylogroup D-38 and D-69 status of Assembly in EngteroBase were also used (Table S1).

### Statistical analyses

Data were indicated as counts, percentages, means, standard deviations (SD), or medians. Student’s t-test or Fisher’s exact test was used to assess numerical and categorical variables. P < 0.05 was considered statistically significant. Data were presented as counts and percentages, means and SDs, or medians and percentiles (25th–75th). Numerical and categorical variables were assessed using Student’s t-test or Fisher’s exact test. Two-tailed P < 0.05 was considered statistically significant. Statistical analyses were performed using GraphPad Prism 8 (GraphPad Software, Inc., La Jolla, CA, USA).

### Ethical considerations

This study was approved by the Ethics Review Board at Kyorin University (#1,696).

Patient identity is secure because of the disclosed details of the patients.

### Nucleotide sequence accession numbers

The sequences of the 23 UPEC strains examined in this study were deposited in the GenBank/EMBL/DDBJ database under accession numbers SAMD00792079 to SAMD00792101.

## RESULTS

### Genotypes, antibiotic resistance, and the set of virulence-associated genes of clinical isolates

Among the 60 bloodstream infections with ESBL-producing *E. coli*, 76.7% were UTIs (46 cases), and the rest were biliary tract infections and gastrointestinal perforations (Table 1). The results of phylogroup analysis by PCR-based methods are summarized in Table 1. Regarding UPEC strains, 93.5% (43 strains) were in phylogroup B2, and the remaining three were in phylogroup D (Table 1). Urosepsis was observed in 8 of 38 cases by ST131 and in 5 of 8 cases by non-ST131 (Table 3). It was significantly less in ST131 than non-ST131 [odds ratio (OR), 0.16; 95% confidence interval (95% CI), 0.038 to 0.873; P = 0.031)]. Regarding antimicrobial agents, all strains were susceptible to imipenem/cilastatin, meropenem, cefmetazole, and latamoxef, whereas the rate of fluoroquinolone nonsusceptibility (R or I) was ∼80.0% for all strains (Table 4). However, the rates of ciprofloxacin and levofloxacin were 89.6% and 87.5%, respectively, for ST131 strains, compared to 75% and 50.0% for non- ST131.

**Table 3.** Characters of patients and genotypes of bacterial strains isolated from UTIs.

| Baseline characteristics of the patients |  | UTIs Total<br>(n=46) | Phylogroup B2 |  | D<br>(n=3) |
| --- | --- | --- | --- | --- | --- |
|  |  |  | ST131<br>(n=38) | non-ST131<br>(n=5) |  |
| Male/Female |  | 15/31 | 14/24 | 1/4 | 0/3 |
| Age (Year) | Median (IQR 25-75) | 84 (72-88) | 86 (78-89) | 79 (68-84) | 59 (57-65) |
|  | Min - Max | 26-96 | 26-96 | 50-95 | 55-71 |
| Underlying diseases | Cancer | 7 | 6 | 1 | 0 |
|  | Diabetes mellitus | 9 | 6 | 2 | 1 |
|  | Immunodeficiency<sup>*1</sup> | 7 | 6 | 1 | 0 |
| Sepsis<sup>*2</sup> |  | 13 | 8 | 3 | 2 |

<sup>\*1</sup>Patients with chronic kidney disease (including patients undergoing dialysis), malnutrition, hypothyroidism, autoimmune diseases, and those receiving immunosuppressive or immunomodulatory agents were classified under immunodeficiency.
<sup>\*2</sup>In accordance with the Third International Consensus Definitions for Sepsis and Septic Shock (Sepsis-3), cases with a Sequential Organ Failure Assessment (SOFA) score of 2 or higher were diagnosed as sepsis.

**Table 4.** Antibiotic susceptibility of clinical isolates.

| Antibiotics | Total (n=46) | ST131 (n=38) | non-ST131(n=8) |
| --- | --- | --- | --- |
| Ampicillin | 46 (100.0%) | 38 (100.0%) | 8 (100.0%) |
| Piperacillin | 46 (100.0%) | 38 (100.0%) | 8 (100.0%) |
| Ceftazidime | 46 (100.0%) | 38 (100.0%) | 8 (100.0%) |
| Cefazolin | 46 (100.0%) | 38 (100.0%) | 8 (100.0%) |
| Cefepime | 46 (100.0%) | 38 (100.0%) | 8 (100.0%) |
| cefmetazole | 0 (0.0%) | 0 (0.0%) | 0 (0.0%) |
| Cefotaxime | 46 (100.0%) | 38 (100.0%) | 8 (100.0%) |
| cefepodoxime proxetil | 46 (100.0%) | 38 (100.0%) | 8 (100.0%) |
| Cefuroxime | 46 (100.0%) | 38 (100.0%) | 8 (100.0%) |
| Latamoxef | 0 (0.0%) | 0 (0.0%) | 0 (0.0%) |
| Aztreonam | 46 (100.0%) | 38 (100.0%) | 8 (100.0%) |
| imipenem/cilastatin | 0 (0.0%) | 0 (0.0%) | 0 (0.0%) |
| Meropenem | 0 (0.0%) | 0 (0.0%) | 0 (0.0%) |
| sulbactam/ampicillin | 35 (76.1%) | 29 (75.0%) | 6 (75.0%) |
| tazobactam/piperacillin | 1 (2.2%) | 1 (4.2%) | 0 (0.0%) |
| Amikacin | 0 (0.0%) | 0 (0.0%) | 0 (0.0%) |
| Gentamicin | 14 (30.4%) | 11 (25.0%) | 3 (37.5%) |
| ciprofloxacin | 39 (84.8%) | 33 (89.6%) | 6 (75.0%) |
| levofloxacin | 36 (78.3%) | 32 (87.5%) | 4 (50.0%) |
| sulfamethoxazole-trimethoprim | 17 (37.0%) | 13 (33.3%) | 4 (50.0%) |

Data are expressed as number and percent (%) of non-susceptible strains to each antibiotic.

### Virulence-associated genes in clinical isolates analyzed by PCR

Multiple factors relating to colonization, iron acquisition, and serum resistance collaborate to express UPEC pathogenicity (12). The representative UPEC virulence-associated genes were chosen and assessed by PCR (Table 5; Table S2). As a result, *csgA*, *fimH*, and *fyuA* were detected in all strains. In contrast, *malX*, *usp*, and *ompT* were detected in all phylogroup B2 strains but not in phylogroup D strains. *traT* and *iutA* were detected in more than half of the B2 strains. However, none of the phylogroup D strains were positive. Overall, significantly fewer virulence-associated genes were detected in phylogroup D strains than in phylogroup B2 strains (Table 5; Table S2). *papAH* and *papGII* or *papGIII*, genes of P fimbriae, also known as P pili, thought to be associated with the development of pyelonephritis (19), were not detected in phylogroup D strains.

**Table 5.** Number of virulent associated genes in clinical isolates analyzed by PCR.

| Virulent associated genes |  | Phylogroup B2 |  | Phylogroup D<br>(n=3) | Total<br>(n=46) |
| --- | --- | --- | --- | --- | --- |
|  |  | ST131<br>(n=38) | non-ST131<br>(n=5) |  |  |
| Type-1 pili | *fimH* | 38 | 5 | 3 | 46 |
| Curli fibers | *csgA* | 38 | 5 | 3 | 46 |
| P fimbriae | *papAH* | 10 | 4 | 0 | 14 |
|  | *papGII* | 9 | 3 | 0 | 14 |
|  | *papGIII* | 1 | 1 | 0 |  |
| Iron utilization | *fyuA* | 38 | 5 | 3 | 46 |
|  | *iutA* | 38 | 4 | 0 | 42 |
| Complement inhibition<br>immune evasion | *kpsMTII* | 35 | 5 | 3 | 43 |
|  | *traT* | 28 | 1 | 0 | 29 |
|  | *ompT* | 38 | 5 | 0 | 43 |
| Pathogenicity island marker | *malX* | 38 | 5 | 0 | 43 |
| Toxin | *cnfI* | 9 | 2 | 0 | 11 |
|  | *hlyA* | 9 | 1 | 0 | 10 |
|  | *usp* | 38 | 5 | 0 | 43 |

The occurrence of virulence-associated genes among ESBL-producing UPEC isolates from bloodstream infections. The sum of *papGII* and *papGIII* positive strains were indicated in the Total column.

### Whole-genome analysis and characterization of clinical isolates

Fifteen were randomly selected from 38 ST131 strains determined by the POT kit and subjected to whole-genome analysis together with eight non-ST131 strains for *in silico* analysis of genotype, serotype, FimH and FumC types, resistance genes, and possession of pathogenic genes (Table 6).

**Table 6.** Characters of the clinical isolates obtained by *in silico* analysis and each patient data.

| Strain | Phylo group | MLS T | O antigen | H antigen | *fimH* type | *fumC* type | CTX-M | Age | M/F<sup>a</sup> | Disease |
| --- | --- | --- | --- | --- | --- | --- | --- | --- | --- | --- |
| KYE006 | D | 38 | O45 | H15 | 24 | 26 | 14 | 55 | F | Pyelonephritis, Septic shock, DIC |
| KYE008 | B2 | 131 | O25 | H4 | 30 | 40 | 27 | 83 | F | UTI, Septic shock |
| KYE011 | D | 38 | O51 | H40 | 5 | 26 | 14 | 71 | F | UTI, Septic shock |
| KYE013 | B2 | 131 | O25 | H4 | 30 | 40 | 27 | 86 | M | Pyelonephritis |
| KYE016 | B2 | 1193 | O75 | H5 | 64 | 14 | 27 | 68 | M | UTI, Septic shock |
| KYE019 | B2 | 131 | O25 | H4 | 30 | 40 | 27 | 64 | F | renal cyst infection |
| KYE020 | D | 38 | O50/O2 | H30 | 5 | 26 | 14 | 59 | F | Pyelonephritis |
| KYE024 | B2 | 131 | O25 | H4 | 30 | 40 | 27 | 65 | F | Renal abscess, Sepsis |
| KYE026 | B2 | 131 | O25 | H4 | 30 | 40 | 15 | 89 | F | UTI |
| KYE027 | B2 | 131 | O25 | H4 | 30 | 40 | 15 | 94 | M | UTI |
| KYE034 | B2 | 95 | O1 | H1 | 16 | 38 | 14 | 95 | F | UTI |
| KYE035 | B2 | 131 | O16 | H5 | 43 | 40 | 27 | 80 | M | UTI, Septic shock |
| KYE040 | B2 | 131 | O25 | H4 | 30 | 40 | 15 | 90 | F | UTI |
| KYE042 | B2 | 73 | O25 | H1 | 12 | 24 | 8 | 84 | F | UTI, Septic shock |
| KYE047 | B2 | 131 | O25 | H4 | 30 | 40 | 15 | 76 | F | Pyelonephritis |
| KYE049 | B2 | 131 | O25 | H4 | 30 | 40 | 15 | 82 | F | Pyelonephritis |
| KYE054 | B2 | 131 | O25 | H4 | 30 | 40 | 15 | 96 | F | Pyelonephritis |
| KYE056 | B2 | 131 | O16 | H5 | 41 | 40 | 27 | 88 | M | acute prostatitis |
| KYE057 | B2 | *<sup>b</sup> | O25 | H4 | 30 | 40 | 27 | 87 | M | UTI |
| KYE059 | B2 | 1193 | no-hit | H5 | 64 | 14 | 27 | 79 | F | UTI, Sepsis |
| KYE060 | B2 | 131 | O25 | H4 | 30 | 40 | 15 | 28 | F | Pyelonephritis |
| KYE062 | B2 | 131 | O25 | H4 | 54 | 40 | 27 | 89 | M | UTI |
| KYE064 | B2 | 1193 | O75 | H5 | 64 | 14 | 27 | 50 | F | Pyelonephritis |

<sup>a</sup>Acronyms F and M stand for Female and Male, respectively.
<sup>b</sup>Single nucleotide substitution (16G→A) was detected in the *adk* allelic profile of ST131.

Genome analysis showed that MLST was consistent with PCR results, except for KYE057, which was determined to be ST131 with the POT kit. KYE057 was defined as a lineage of ST131, which has a single nucleotide substitution (16G→A) in the *adk* gene detected to ST131 and was included in ST131 strains for statistical analysis in this study. Non-ST131 strains were determined as ST95, ST73, or ST1193 in phylogroup B2 and ST38 in phylogroup D. As in previous results (20), the ST1193 strain showed a disruption of *lacY* by frameshift mutation (Table S3). As for the O antigen type, O25 was the most common (14 strains), followed by O16, O1, and O75, frequently detected in UPEC. ESBL genotypes were CTX-M27 (11 strains), CTX-M15 (7 strains), CTX-M14 (4 strains), and CTX-M8 (1 strain). ST131-O25:H4 H30 pandemic clones were CTX-M15 or CTC-M27, and three ST38 strains were CTX-M14 (Table 6).

### Virulence-associated genes in clinical isolates analyzed *in silico*

Based on virulence gene results by ABRicate (Table S4), ST131 clinical isolates revealed the equivalent number of virulence-associated genes, including the UPEC ST131 type strain, EC958, and they were significantly fewer than the non-ST131 strains (<0.0001; Table 7). Results obtained by PCR were consistent with those of *in silico* data. Well-known UPEC-specific genes, including *ompT*, *usp*, and *malX*, were detected in all phylogroup B2 strains *in silico* analysis in addition to PCR. In contrast, they were absent in ST38 strains.

**Table 7.** Number of virulent associated genes analyzed *in silico* (n=23)

| Phylogroup | MLST | Median (IQR 25-75) | Min - Max |
| --- | --- | --- | --- |
| B2 | ST131 (n=15) | 192 (190 - 200) | 174 - 207 |
|  | non-ST131(n=5) | 220 (219 - 228) | 209 - 233 |
| D | non-ST131 (n=3) | 227 (223 - 234) | 218 - 240 |

Data are expressed as median (IQR 25-75), minimum (Min) and maximum (Max) number of virulence-associated genes detected by ABRicate were indicated. The virulence-associated genes in 536, CFT073, CFT073, UTI89, and EC958 were 261, 234, 257 and 198, respectively. The number of virulence-associated genes was tailed from the numbers of genes indicated in Table S4.

Clustering analysis of clinical isolates, including UPEC reference strains, was performed based on virulence genes. As a result, each clade consisted of strains dependent on the genotype (Fig. 1A). Focusing on the MLST-specific genes, the genes found only in ST38 clinical isolates included genes of *epa* (the members of secondary T3SS-*epa*), *esp* (EPEC- T3SS secreted protein, T3SS-*esp*), *ycb*, and the *hlyE* gene (Fig. 1B; Table S4). Most of ST131 and all ST95 and ST73 and ST1193 specific genes were hypothetical genes (Fig. 1B; Table S4).

**Fig. 1.**
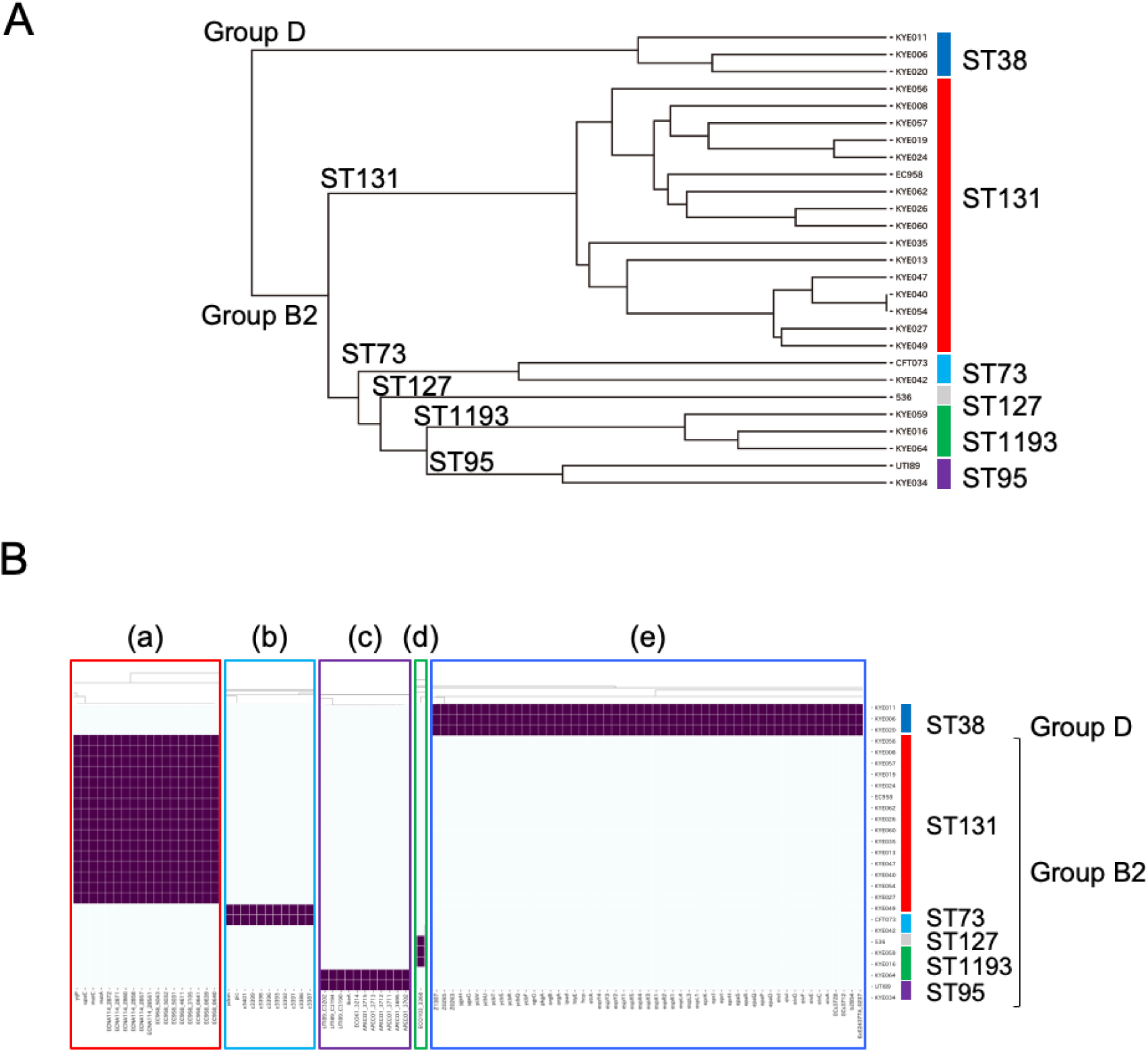
The ESBL-producing UPEC clinical isolates and representative UPEC strains clustered according to virulence-associated gene carriage detected by genome analysis. The presence of the gene is indicated with purple. Dendrograms of bacterial strains were deposited to distinguish the results (A). The genes specifically detected in the ST131 (a), ST73 (b), ST95 (c), ST1193 (d) and ST38 (e) strains were shown (B). Phylogroups B2 and D are indicated with Group B2 and D.

### Homologs of virulence-associated genes and gene loci relating to their functions

The hypothetical genes found in MLST-specific genes were analyzed by BLAST search using the amino acid sequences and gene order in Prokka. Accordingly, many genes were annotated as T6SS gene members and found upstream and downstream of c3400, present in ST131 and other phylogroup B2 strains (Table S5). Based on the nucleotide sequences, it was shown that c3400 encoded *tssF* and *tssG* of T6SS-1. T6SS is classified into three distinct groups, T6SS-1 to -3, according to genetic structures. Other T6SS-1 gene homologs were consecutively found around the c3400 genes in other phylogroup B2 strains. Furthermore, utilization of the homologs is likely to be MLST-dependent (Table S5). In contrast, c3400 and any of the T6SS-1 homologs were not found in ST38 strain genomes of ST38 and ST69 strains in EnteroBase (data not shown).

Further investigation was performed to find the other gene homologs of hypothetical genes detected using BLAST search, and the order of genes obtained by Prokka analysis was referred to find the other gene homologs of hypothetical genes (Tables S5 and S6).

Based on ABRicate, Prokka, and BLAST search results, gene homologs or genes designated with several names were analyzed and annotated (Table S5). Then, the genes containing in same operon or functional genes loci were grouped (Table S7). Based on gene loci (Table S7–S9), clustering analysis was performed, including the four well-known specific virulence genes (*cnf1*, *ompT*, *traT*, and *usp*; Table S9).

Gene loci for the attachment and colonization [type I pili, curli biosynthesis, F9 fimbriae, PpdD pili, T4a pili, *E. coli* common pilus], the iron acquisition systems [enterobactin, yersiniabactin, hemin, and Sit iron/manganese utilization system], and motility and chemotaxis, and Group 2 capsule were found in all strains, except for KYE013 (Fig. 2; Table 8). Fewer secretion systems were detected in ST131 strains than in non-ST131 strains (Table 9).

**Fig. 2.**
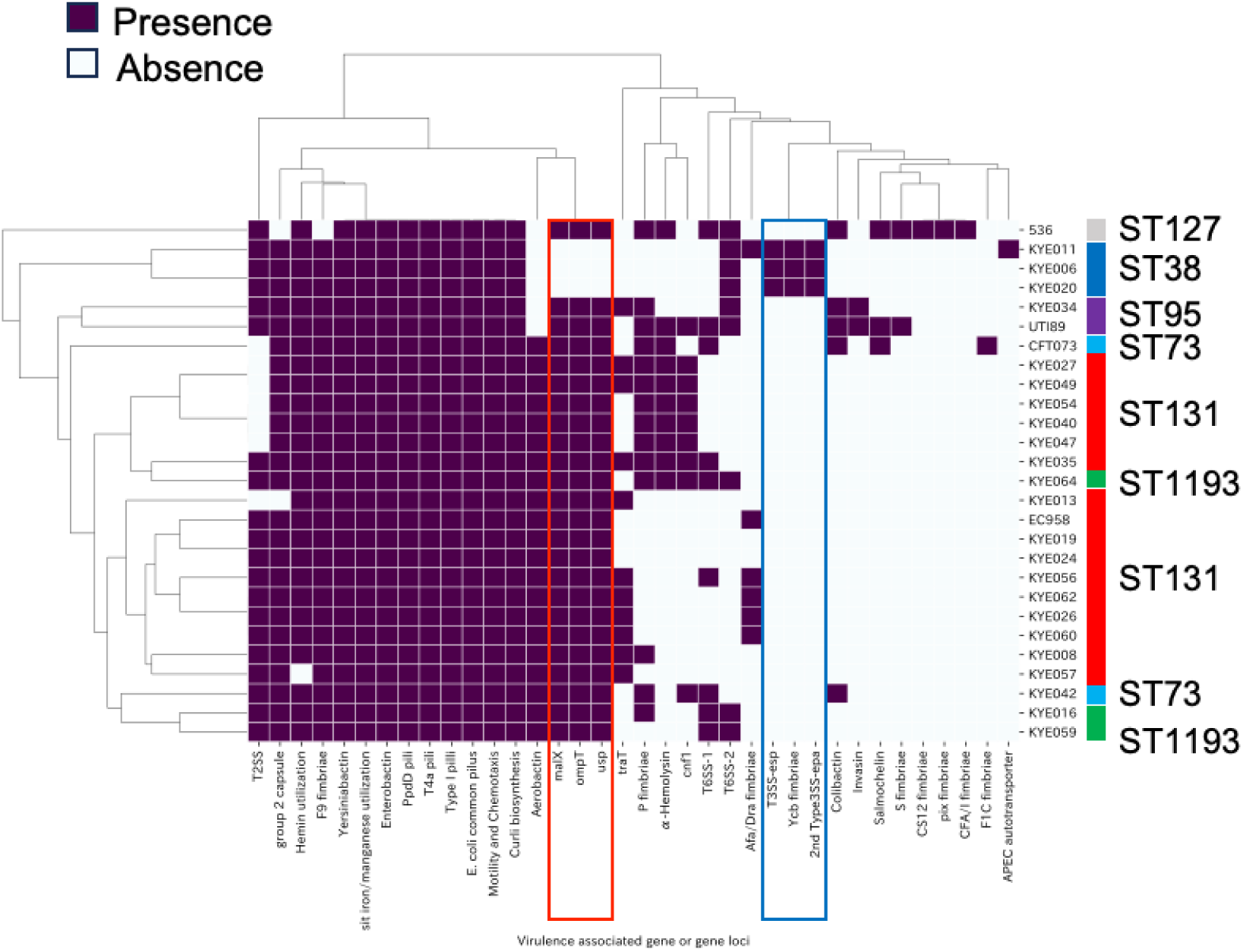
Hierarchical clustering of the bacteria based on the virulence-associated gene loci and genes profiles. The gene loci containing the predicted functional gene sets showing purple were determined (Table S7 - S9). The genes that were specifically not detected in ST38 strains were indicated with a red box. The gene loci that were specifically detected in ST38 strains were indicated with a blue box.

**Table 8.** Presence and absence of virulence-associated gene or gene loci.

| Classification | Gene loci |  |
| --- | --- | --- |
| Gene loci commonly detected in the ESBL-producing UPEC clinical isolates | Fimbriae | Type I pili<br>Curli biosynthesis<br>F9 fimbriae<br>PpdD pili<br>T4a pili<br>*E coli* common pilus |
|  | Iron utilization | Enterobactin<br>Yersiniabactin<br>Sit iron/manganese utilization<br>(Hemin utilization) |
|  | Motility | Motility and Chemotaxis |
|  | Immune evasion | (group 2 Capsule) |
| phylogroup D-ST38 specific gene or gene loci | Fimbriae | Ycb fimbriae |
|  | Secretion system | 2nd T3SS-*epa*<br>T3SS-*esp* |
| Gene or gene loci that were not detected in phylogroup D-ST38 specifically<br>phylogroup D-ST38 specific gene or gene loci | UPEC PI marker | *malX* |
|  | Serum resistance | *ompT* |
|  | Fimbriae | Ycb fimbriae |

The virulence-associated genes were grouped and indicated as gene loci according to the Table S7.
The gene loci found in all clinical isolates except for one strain were parenthesized.

**Table 9.** Number of virulence genes gene loci detected in the clinical isolates examined.

|  | Phylogroup B2 |  |  |  | Phylogroup D |  |
| --- | --- | --- | --- | --- | --- | --- |
|  | ST131 |  | non-ST131 |  | ST38 |  |
|  | Average | (Min-Max) | Average | (Min-Max) | Average | (Min-Max) |
| Adhesin (9) | 6.7 | (6-7) | 6.8 | (6-7) | 7.3 | (7-8) |
| Fe acquisition system (5) | 4.9 | (4-5) | 4.8 | (4-5) | 4.0 | (4-4) |
| Secretion System (5) | 0.7 | (0-2) | 2.4 | (1-3) | 4.0 | (4-4) |
| Others (6) | 2.3 | (1-3) | 2.8 | (2-4) | 2.3 | (2-3) |

Data are expressed as average, and minimum (Min), maximum (Max) number of gene loci consisting of functional gene members.

The T6SS apparatus requires at least 13 genes to construct (21). Therefore, the secretory apparatus of T6SS-1 predictably lacked ST131-O25 strains and KYE034 (ST95), as reported previously (22) (Fig. 2; Tables S5 and S8-9).

*aec* genes were predicted as members of T6SS-2 (Table S5). There were no *aec* genes in ST131 clinical isolates and the genomes of 51 strains in EnteroBase. Other T6SS-2 gene member homologs were found in all strains, including ST38, except for ST131 strains (Table S5). Since most T6SS-2 genes were lacking in previous data (21), ST73 strains, KYE042, and CFT073 would not have functional T6SS-2. Namely, functional T6SS-2 loci were lacking in ST131 and ST73 strains (Fig. 2; Tables S5 and S8-9). Like T6SS-1 genes, MSLT dependence upon homolog utilization was observed in T6SS-2 genes (Table S5).

P fimbria was encoded by 11 *pap* (pyelonephritis-associated pili) genes (Table S7) linked to pyelonephritis, in addition to UTIs (19, 23, 24). Whole-genome analysis revealed that none of the *pap* genes were present in ST38 strains and the disruption of *pap* operon by the *IS* insertion downstream of *papI* in several phylogroup B2 strains, which were negative for *papA* and *papG* genes by PCR (Tables S2 and S8). Ten of 38 ST131 isolates indicated *papAH* and *papG* positive, indicating they could probably express a functional P fimbriae. This ratio was significantly lower than the four of five isolates with non-ST131 (10 of 38; OR, 0.089; 95% CI, 0.0071–0.7031; P = 0.032).

Functional T3SS-*esp*, 2nd T3SS-*epa*, Ycb fimbriae genes, and *hlyE* were found only in ST38 strains (Fig. 2; Table 8 and 9). ST69 belongs to phylogroup D and is also frequently detected in UPEC strains. The presence of *ycb* genes was assessed using the genome of seven of ST38 strains and 161 of ST69 strains isolated from UTIs in EnteroBase (Table S1).

Accordingly, *ycb* genes were detected in all ST38 strains but not in the ST69 strains examined. Contrarily, 2nd T3SS-*epa*, T3SS-*esp* genes, and *hlyE* were detected in all ST38 and >90% in ST69 strains (data not shown).

## DISCUSSION

This study analyzed the genotype and virulence-associated genes in ESBL-producing UPEC clinical isolates from the bloodstream to elucidate the repertories of virulence- associated factor(s) causing invasive UTIs.

Moreover, 82.6% of clinical isolates indicated ST131, the most prevalent genotype in ESBL-producing UPEC strains consisting of typical ExPEC responsible for bloodstream infections (15, 25). The rest of phylogroup B2 consisted of ST73 and ST95, called “classic STs,” which are common in ExPEC, colonize the intestine, and are frequently isolated from UTIs and bloodstream infections (26) and ST1193, which is considered an emerging clone (27) and is related to the outbreak clone after ST131 lineage (20, 28). ST38 in phylogroup D is among the top 10 human pandemic lineages among ExPEC (29). However, ST38 ExPEC strains have been poorly investigated, except for the comprehensive phylogenomic analysis by Chowdhury et al. (25).

Because the resistance rate was ∼10%, fluoroquinolones have been recommended to treat UTIs, including ESBL-producing *E. coli* UTI cases, based on the clinical outcomes of the *E. coli* ST131 clonal group (30). However, ∼80% of the strains isolated from UTIs and ∼90% of ST131 strains were nonsusceptible (Table 4). Although the possibility of the bias used by past antimicrobials cannot be excluded because of bloodstream infections, the results might indicate that treatment is becoming more refractory.

When analyzing representative UPEC virulence-associated genes, the design PCR primers were repeated twice or thrice for *kpsMT*, *papAH*, and *papG* genes based on the homologous region of the genes of open genomes (data not shown). Primers should be carefully designed considering the polymorphism frequently observed in genes encoding bacterial surface proteins. Eventually, only three genes were detected in all strains examined, and nine genes were not detected in ST38 strains by PCR (Table 5; Table S2). Consequently, less virulence-associated genes were detected in ST38 strains by PCR (Table 5; Table S2). Contrary to PCR results, whole-genome analysis revealed a smaller number of virulence-associated genes in ST131 strains than in non-ST131 strains (Table 7; Table S4). Furthermore, there was dependence of virulence-associated genes on MLST or, more notably, on phylogroup (Fig. 1A).

*ycb* genes were detected prominently in ST38 strains but not in phylogroup B2 clinical isolates (Fig. 1B; Table S4, S8) and in the genomes of ST69 strains in EnteroBase (data not shown). The *ycb* operon was previously detected in enteropathogenic *E. coli* strains but is present in a cryptic operon system in a non-pathogenic *E. coli* strain, K12 (31, 32) and related to the invasion of the human ileocecal epithelial cell line HCT-8 (32). Therefore, *ycb* genes could be associated with the virulence of ST38 strains. Further analyses will elucidate the expression and functions of Ycb in ST38 strains.

Although *IS* disrupted the locus in several B2 strains, the P fimbriae gene locus was detected in all phylogroup B2 strains. The *papGII* locus, which is considerably associated with invasive infections and severe UTIs (24, 33). Furthemore, the *papA* gene was present in 75% of ST69 strains in EnteroBase (data not shown). It was considered possible that some of the *pap* genes may have been lost during evolution or that genes deficient in *pap* may have been transmitted horizontally in these strains. On the other hand, none of the ST38 strains had any of the *pap* gene loci.

By predicting the functions of hypothetical genes in clinical isolates, T6SS-1 gene homologs around the c3400 genes were detected in all phylogroup B2 strains but not in ST38 strains (Table S4, S5). T6SS is a secretion system derived from bacteriophages and is found in Gram-negative bacteria, frequently found in pathogenic strains and the absence of nonpathogenic strains (22). The nature of T6SS-1 effector proteins is different. Phenotypes and effectors associated with T6SS-1 are enteroaggregative *E. coli* (EAEC) and avian pathogenic *E. coli* (21). Similar to the results of avian pathogenic *E. coli* by Tantoso *et al*. (34), T6SS-1 genes were absent in phylogroup D strains, and several essential T6SS-1 genes were lacking in ST131-O25:H4 clones and KYE034 (ST95; Table S8).

T6SS-2 genes were found in all strains, except for ST131, although most were lacking in ST73 strains (Table S8). *E. coli* T6SS-2 was initially found in EAEC. T6SS-2 would be associated with phylogroups D to F but not B2 (35). However, T6SS-2 was detected in ST95 in this study and elsewhere (34). According to Prokka analysis, genes encoding putative Rhs-family proteins predicted T6SS-2 effectors were found upstream of the *vgrG* gene in ST38 stains (data not shown) (36). Therefore, ST38 strains might utilize different effectors from other phylogroup B2 strains. The biological approach will unveil the function and role of T6SS in *E. coli* pathogenicity.

Regarding the absence of P fimbriae genes, *usp*, *ompT*, and *malX*, UPEC-specific genes, and the presence of multiple EPEC genes, such as Ycb, 2nd T3SS-*epa*, and T3SS-*esp* loci genes, ST38 strains are different from other UPEC strains (Table 8). UPEC causes infection ectopically from the intestinal tract to the urinary tract and may have shared more genes with other pathogenic *E. coli* than previously thought. Hybrid strains that share genes characteristic of EPEC and UPEC have emerged in recent years (37). ST38 NDM-5-producing *E. coli* isolates caused an outbreak in the Czech Republic (38). Genomic characterization and pathogenicity of ST38 clinical isolates are required to monitor future trends.

Type I pili, *E. coli* common pilus, PpdD pili, T4a pili, Curli biosynthesis, F9 fimbriae, enterobactin, yersiniabactin, Sti iron/manganese utilization systems, and motility genes were found in all clinical isolates, and Group 2 capsule and hemin utilization gene loci were found in all strains but one examined in this study (Table 8). These factors have a crucial role in the pathogenesis of bloodstream infections by UPEC.

Although a limited number of patients were analyzed in this study, sepsis cases were detected less in ST131 (Table 3). However, less virulence-associated gene results with PCR were obtained in ST38, non-ST131 strains, in previous results using the VFDB database for ABRicate. Therefore, *E. coli*_VF was adopted for the database to increase the virulence-associated gene target. Consequently, the number of virulence-associated genes especially secretion system genes was significantly lower in ST131 strains than those in non-ST131 strains (Table 7 and 9). ST131 strains were recognized as highly virulent UPEC clones (33). In contrast, ST131 isolates do not consistently express higher virulence potential compared to different *E. coli* types causing invasive extraintestinal infections using the mouse sepsis model (39). The correlation between severity and virulence factors is not clear (40). However, since the probably related to *E. coli* virulence were detected less in ST131 strains, ST131 clones might have become attenuate during spreading infections.

Increasing the detection rate of ESBL-producing *E. coli* will make treating severe UTIs, such as pyelonephritis and urosepsis, more difficult (41, 42). Prediction of the virulence of ESBL-producing *E. coli* strains is more critical for diagnosis and treatment. With further enhanced *in silico* analysis, UPEC virulence gene and clinical information will unveil ESBL-producing UPEC pathogenicity.

## Acknowledgments

We thank T. Nozaki for the technical assistance in laboratory work. We thank Prof. T. Sato for the assistance with the statistical analysis in this research. Their expertise and support were instrumental in the successful completion of this study. This work was supported by JSPS KAKENHI grant nos. 22K09535 and 24K11640.

